# Reconstructing the blood metabolome and genotype using long-range chromatin interactions

**DOI:** 10.1101/656132

**Authors:** Tayaza Fadason, William Schierding, Nikolai Kolbenev, Jiamou Liu, John Ingram, Justin M. O’Sullivan

## Abstract

The mechanisms of metabolism comprise a large number of biochemical pathways with a myriad of poorly characterised genetic influences. In this study, we perform a systematic integration of chromatin interaction (Hi-C), expression quantitative trait loci (eQTL), gene ontology, drug interaction, and literature-supported connections to deconvolute the genetic regulatory influences of 145 blood metabolite-associated single nucleotide polymorphisms (SNPs). We identify 577 genes that are regulated via chromatin looping to 130 distal and proximal SNPs across 48 different human tissues. The affected genes are enriched in categories that include metabolism, enzymes, plasma proteins, disease development, and potential drug targets. These novel SNP-gene-metabolite associations are a valuable resource for understanding the molecular mechanisms guiding pathologic metabolite levels in human tissues, and for further investigation into disease diagnosis and therapy.

## Introduction

The interaction between genetic variation, environment and lifestyle affects metabolite levels in humans^1^. The levels of these metabolites are central to the expression of phenotypes and disease development *e.g.* cholesterolaemia, cancer, and type 2 diabetes^2–4^. Despite decades of work targeting metabolites of interest, the genetic networks and genes that modify metabolic potential remain poorly characterised and their therapeutic and diagnostic utility remains unrealized.

Whole genome or exome sequencing, in combination with metabolomic techniques, have been used to identify the genetic influencers of the human blood metabolome^5–9^. In arguably the most comprehensive analysis to date, Shin and colleagues^5^ performed a genome-wide association study (GWAS) and identified 145 genetic loci (SNPs) that correlate with the levels of approximately 400 blood metabolites. As is typical of all GWAS studies, >65% of the SNPs identified by Shin *et al*. were in non-coding regions of the genome. Shin *et al*. predicted causal genes of the genetic loci by scanning 500 kilobase (Kb) regions flanking the SNPs for genes whose functions linked with the corresponding metabolite. While it is commonly accepted that GWAS SNPs are enriched in regulatory elements, identifying the genes affected by the SNPs using a nearest-relevant-gene approach is problematic because the accumulating evidence supports long-range gene regulatory interactions, including inter-chromosomal, between regulatory elements and genes^10–13^. Moreover, recent studies suggest that long-range regulatory interactions may play a more significant role in modifying disease outcomes than proximal interactions^14–16^. Long-range gene regulatory interactions can occur by several mechanisms: 1) epistatic interactions between gene products; 2) widespread action of a transcription factor or non-coding RNA; 3) scanning (*e.g.* the spreading of silencing complexes along a chromosome); or 4) chromatin looping (*e.g.* direct physical contact between enhancers and gene promoters)^17^. We, and others, have previously used genome structural data captured by proximity ligation techniques (*e.g.* Hi-C^18^) and eQTL data (*e.g.* GTEx^19^) to detect chromatin loops that bring long-range regulatory loci in spatial proximity with their target genes^14,20–24^.

Here, we set out to use the 3D genome organization to identify target genes for the 145 blood metabolite-associated SNPs reported by Shin *et al*.^5^. To achieve this, we interrogated chromatin interaction (Hi-C) data from eight human cell lines^25^ (Supplementary Table 1) to identify genes that are in physical contact with the SNPs^24^. Next, we designated functional SNP-gene associations by interrogating expression quantitative trait loci (eQTL) in 48 human tissues from the Genotype-Tissue Expression (GTEx^19^, www.gtexportal.org) database^24^. Finally, we employed an array of approaches including gene ontology (KEGG, www.genome.jp), protein classification (Protein Atlas, www.proteinatlas.org), drug target analysis (DGIdb, www.dgidb.org), and literature text mining (PubMed®, www.ncbi.nlm.nih.gov/pubmed) to annotate the genes that are involved in modulating intermediate metabolites in human blood (Supplementary Figure 1).

**Figure 1.**
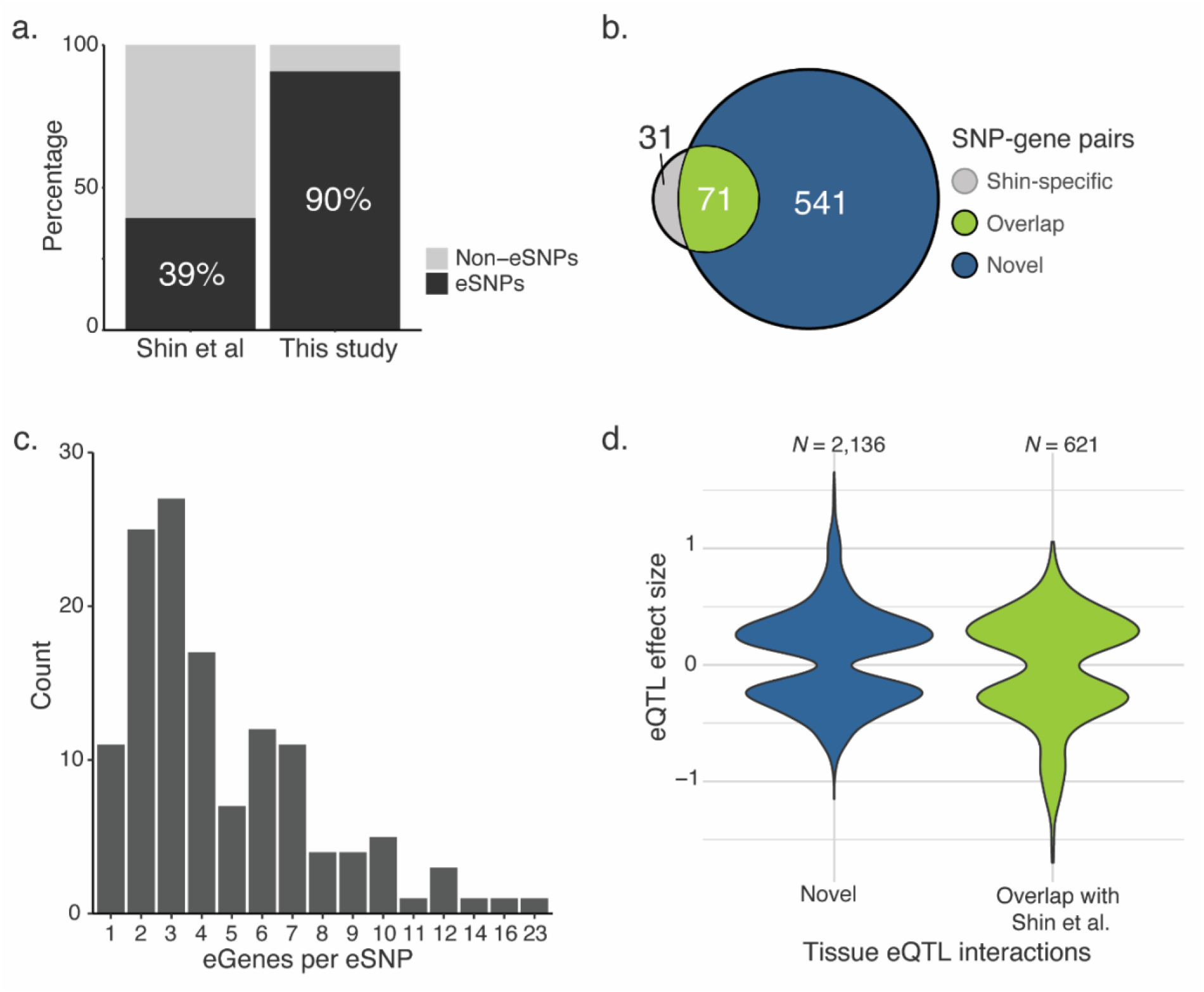
Regulatory interactions involving metabolite-associated SNPs. a) 90% of metabolite-associated SNPs mark eQTLs involved in long-range regulatory interactions. b) 71 of 102 of the causal SNP-gene pairs predicted in Shin et al are among the 612 eSNP-eGene pairs we identified. c) Bar plot of number of genes regulated per eSNP (mode = 3). d) Violin plots of the normalized effect sizes (beta) of the eQTL interactions in different tissues.

## Results

### Metabolite-associated SNPs mark eQTLs

Shin *et al*. reported 145 metabolite-associated loci that are marked by SNPs with genome-wide significance. 102 of these loci were assigned predicted causal genes based on proximity to the SNPs and an established association between the gene and metabolite^5^. Notably, only 57 (∼39%) of the loci were reported as eQTLs by Shin *et al* (Figure 1a). Integrating genomic organization into the analysis identified eQTLs for 130 (∼90%) of the 145 metabolite-associated SNPs (hereafter eSNPs), more than double that reported by Shin *et al*. (Figure 1a, Supplementary Table 2). None of the 15 non-eQTL SNPs (Supplementary Table 3) in our study was reported as being involved in an eQTL by Shin *et al*. This is consistent with a different mode of action, or specificity in the developmental timing of the linkage of these SNPs to their metabolites.

The 130 eSNPs we identified were associated with the expression of 577 genes (*i.e.* eGenes) through 612 unique eSNP-eGene pairs (Figure 1b). The eSNP-eGene pairs included 68.6% of the causal SNP-gene pairs predicted by Shin et al (Figure 1b, Supplementary Table 4). Notably, the eSNPs were associated with between 1 and 23 eGenes, with 3 eGenes as the mode (Figure 1c), in a tissue specific manner. This is consistent with previous reports on shared gene regulatory sites^24,26,27^. Altogether, we identified 2,757 eSNP-eGene interactions in 48 different human tissues (Supplementary Table 5) of which 621 overlapped Shin *et al*.’s predictions (effect size range, -1.69778 to 1.05834), while 2,136 are novel (effect size range, -1.15093 to 1.65858; Figure 1d).

### Metabolite-associated SNPs target genes in metabolic pathways

We annotated the eGenes’ biochemical functions using the Kyoto Encyclopedia of Genes and Genomes (KEGG) Pathway database. 218 (37.8%) of the eGenes are annotated as being involved in ≥1 biochemical pathway (Supplementary Table 4). As expected, the largest single set of annotations for the eGenes is metabolism (*n* = 95, Figure 2a). The next most represented categories *i.e.* organismal systems (*n* = 75) and human diseases (*n* = 65) each share 12 genes with the metabolism category (Figure 2a). Classification of the eGenes using The Human Protein Atlas identified significant enrichment (bootstrapping of null sets) of enzymes, plasma proteins, disease related genes and potential drug targets within 490 of the eGenes (Figure 2b, Supplementary Table 4). The observed enrichment in the metabolic pathway and protein classification analyses is consistent with the eGene products acting as modifiers of metabolic activity.

**Figure 2.**
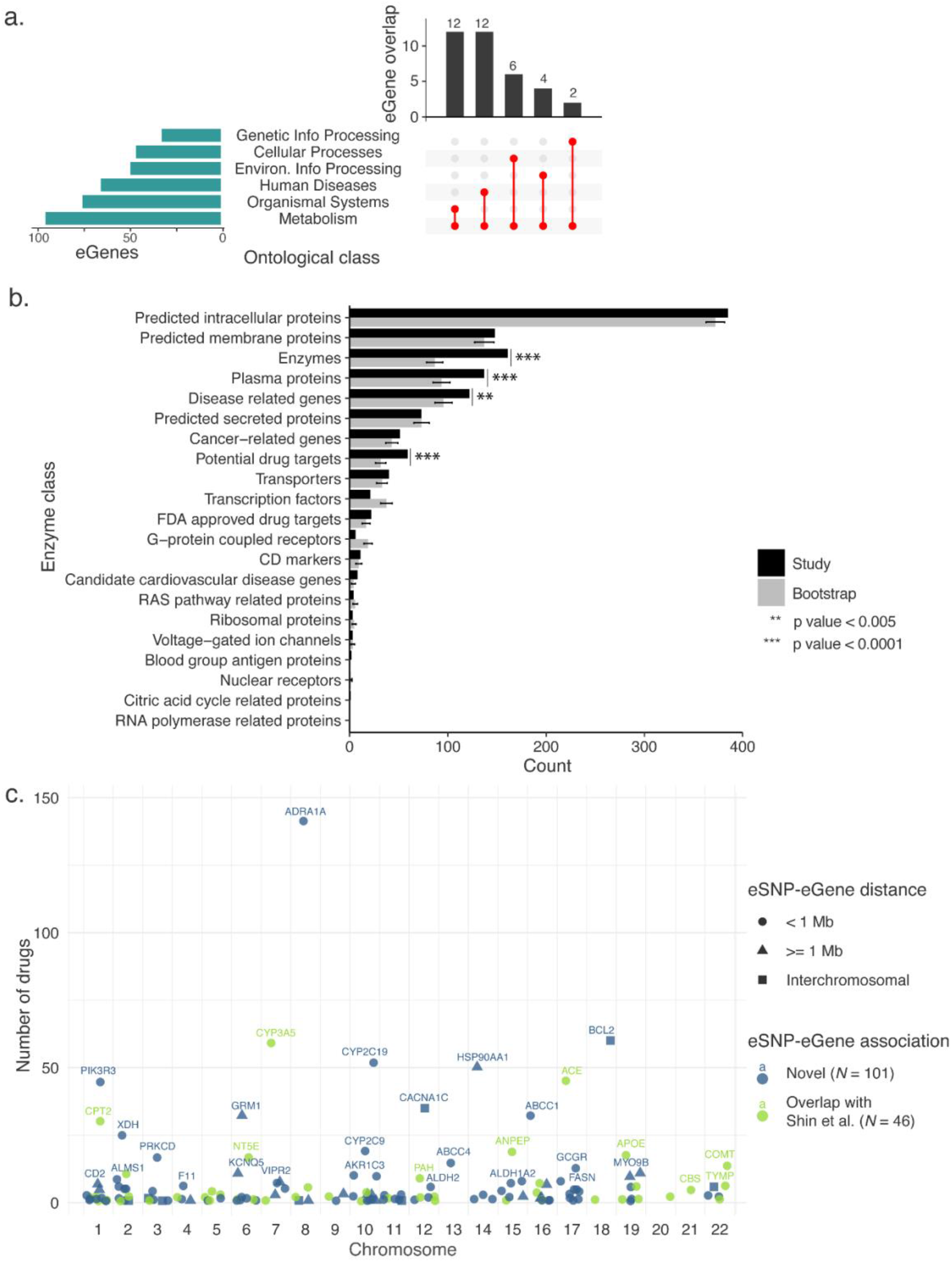
Functional annotation of eGenes associated with metabolite eSNPs. a) eGenes are enriched for metabolism (horizontal bars) in KEGG pathways analysis. The metabolism genes are also involved in other pathways (vertical bars). b) Protein classification of genes is significantly enriched for enzymes, plasma proteins, potential drug targets, disease related genes. Error bars represent one standard deviation of 10,000 bootstraps. c) 147 eGene products are druggable based on DGIdb analysis. 68.7% (101) of the druggable products are encoded by eGenes that were not previously linked to the metabolite-associated SNPs.

We screened the Drug Gene Interaction database (DGIdb) to identify which of the eGenes encode potentially targetable products. Products of 147 (25.5%) of the eGenes are targets of at least one drug (Figure 2c, Supplementary Table 6). 68, 119, and 142 of the druggable eGenes are also involved with metabolism, >1 biochemical pathway, and >1 protein class respectively. 101 of the 147 druggable eGenes have not been previously linked to genetic variants associated with metabolism.

There were several notable examples of metabolite associated gene–eSNP networks. For example, *APOE*, which has been associated with modulation of total cholesterol^34^, was reported by Shin *et al*. as an eGene and predicted as the causal gene of nearby (3 Kb) cholesterol-associated rs445925. In addition to *APOE* (in suprapubic skin), our study revealed novel eQTL for rs445925 with *BCAM* (in basal ganglia, 90 Kb away), *RYRI* (in skeletal muscle, 6.3 Mb away), and *RERE* (in suprapubic skin, on chromosome 1) (Figure 3a).

**Figure 3.**
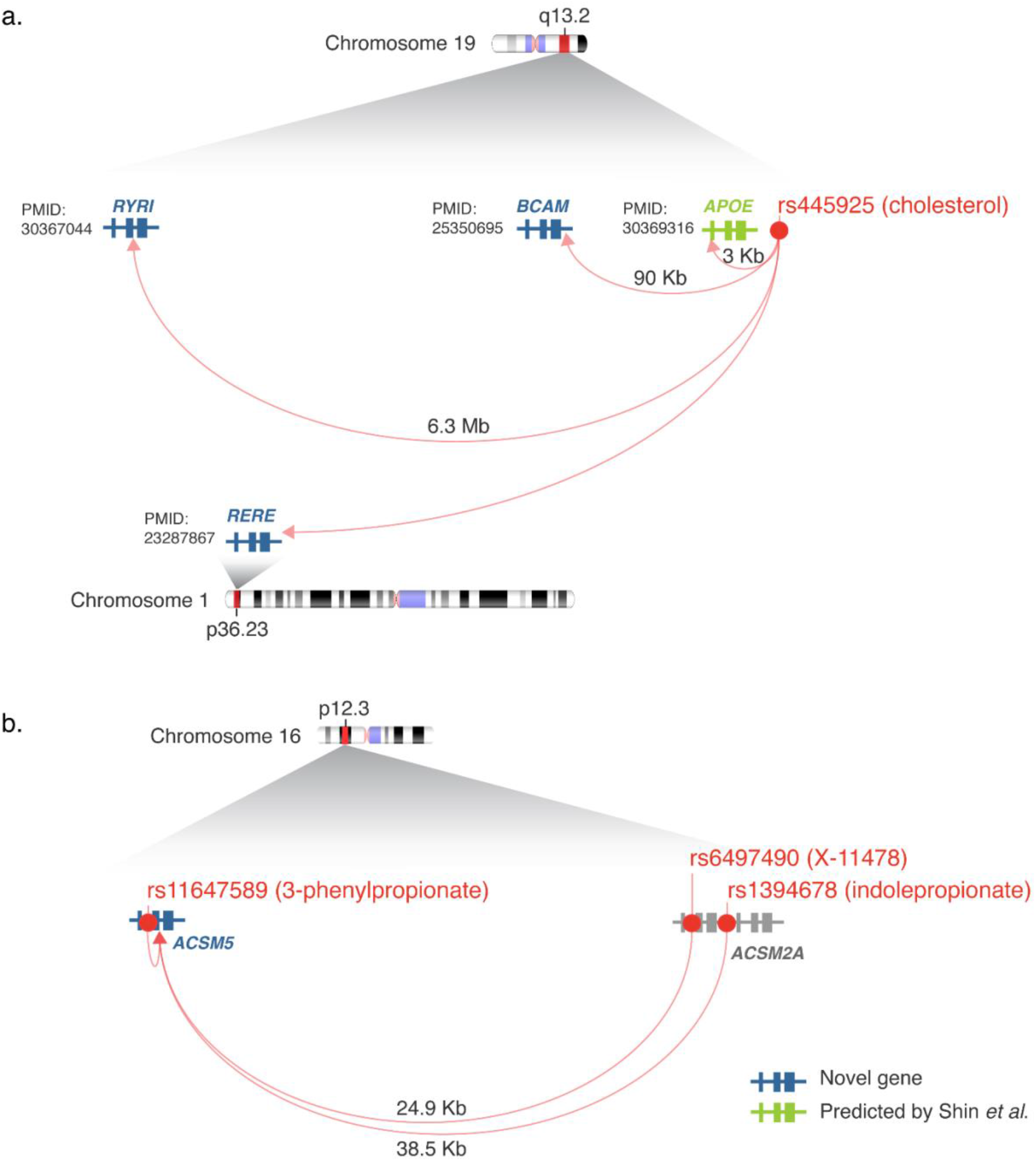
eGene-metabolite associations. a) Cholesterol-associated rs445925 marks an eQTL that correlates with expression levels of *APOE, BCAM, RYRI*, and *RERE*. All four genes have literature (PMIDs)28–31 supporting their links with cholesterol. b) *ACSM5* pairs as an eGene with rs11647589, rs6497490, and rs1394678, which were associated with 3-phenylpropionate, X-11478 (an unknown metabolite), and indolepropionate respectively. The metabolism of 3-phenylpropionate and indolepropionate requires a Coenzyme A ligase such as *ACSM5*32,33.

We also identified network examples where eGenes linked to multiple regulatory hubs. For example, Acyl-CoA Synthetase Medium Chain Family Member 5 (*ACSM5*) associates with eQTL SNPs rs11647589 (in 16 tissues, intronic in *ACSM5*), rs1394678 (in 14 tissues, 38.5 Kb from ACSM5, intronic in *ACSM2A*), and rs6497490 (in tibial artery, 24.9 Kb from ACSM5, intronic in *ACSM2A*). rs11647589, rs1394678, and rs6497490 are associated with 3-phenylpropionate, indolepropionate, and X-117478 (an unknown metabolite) respectively (Figure 3b). rs11647589, rs1394678 and rs6497490 mark distinct eQTLs because they are not in linkage disequilibrium (defined as LD >= 0.8 in the EUR population, HaploReg v4.1^35^).

### Literature text mining supports eGene-metabolite associations

We conducted a stringent semi-automated literature text mining to identify literature that supported a link between the eGenes and the metabolites with which the eSNPs are associated. Firstly, we queried the MEDLINE® database using PubMed® APIs for research article titles or abstracts that contain the eSNP-eGene, eSNP-metabolite, or eGene-metabolite pairs (Figure 4a). We then manually curated articles supporting eGene-metabolite pairs. The manual curation confirmed literature support for 41 eGene-metabolite pairs, 14 of which were not reported by Shin *et al*. (Supplementary Table 7). 29 of the literature-supported associations include genes that are involved at least one biochemical pathway and whose products are druggable (Figure 4b and c).

**Figure 4.**
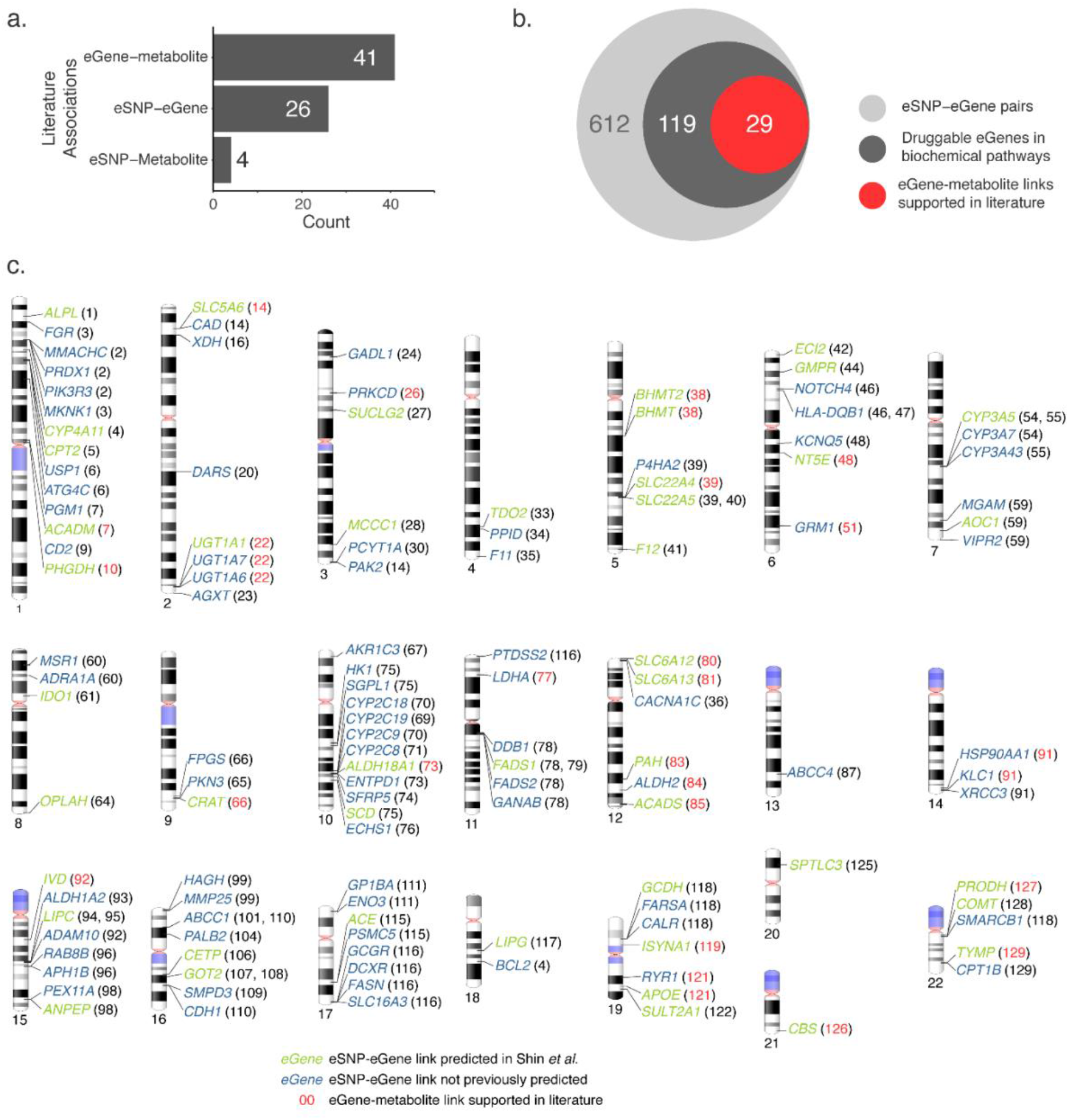
Functional annotation of eGenes associated with metabolite eSNPs. a) There is more literature support for eGene-metabolite associations than for eSNP-eGene or eGene-metabolite associations. b) A schematic of eGenes that are in biochemical pathways, encode druggable products, and whose relationship with their SNP metabolite has literature support. c) Ideogram of 119 druggable eGenes in b. Genes are colored green if the SNP-gene association was reported in Shin et al., or blue if it is not. Numbers in brackets correspond to the locus number (in Supplementary Table 2) of the eSNPs that spatially interact with the eGenes. The red-colored numbers indicate that the relationship between an eGenes and its SNP-associated metabolite (Supplementary Table 4) is supported in literature via text mining.

The literature supported the putative functional outcomes for the rs445925-APOE, BCAM, RYR1, RERE network as follows: 1) *BCAM* had a suggestive association in a GWAS meta-analysis of LDL cholesterol response to statins^29^; 2) *RYRI*, together with other nonalcoholic steatohepatitis related genes, has been linked to dietary cholesterol^30^; and 3) *RERE* was associated with circulating blood CD34+, which positively relates to total cholesterol^31^. Therefore, we contend that these eQTL associations not only link SNPs to genes that are relevant to cholesterol, they also reveal the tissues that might be important for its metabolism.

The absence of literature support for eGene-metabolite associations does not equate to an absence of relationship. For example, the intergenic rs7809615 is associated with levels of androsterone sulfate and 4-androsten-3beta,17beta-diol disulfate 2 in the blood metabolite GWAS (Supplementary Table 2). Our method, as well as that of Shin et al., links rs7809615 to *CYP3A5* (Figure 4c), which encodes a cytochrome P450 known to be active against steroids such as testosterone^36^. rs7809615 also links with *CYP3A7* (Figure 4c), which encodes another P450. Although our stringent text mining did not find articles that directly link the two target eGenes, the STRING database (v10.5, accessed 26/04/2019) reports a binding interaction between *CYP3A5* and *CYP3A5*.

## Discussion

We systematically integrated evidence from chromatin interactions, eQTL data, gene ontology, protein classifications, drug interactions, and the published literature to identify and characterize genes that are regulated by metabolite-associated SNPs. Our results provide a step-change in our current understanding of the variant-gene regulatory interactions of metabolism, particularly those that rely on chromatin looping.

The integration of chromatin interaction and eQTL data in this study enabled the mapping of long-range regulatory interactions for ∼90% of the metabolite-associated SNPs. This mapping was robust to the different data sources that were used to identify the eQTLs (GTEx^19^) and the blood metabolite associated SNPs (KORA and Twins UK)^5^. The identification of associations between metabolite-associated eSNPs and eGenes in non-metabolism ontological classes, or with unknown functions provides novel data-driven avenues for empirical testing to delineate the roles played by these eGenes in metabolism. Similarly, the eSNP-eGene connections reported here for variants associated with unknown metabolites could aid the identification of unknown metabolites.

The limited literature support, identified by text mining, for the eGene-metabolite associations can be attributed to: a) the stringent condition that both terms must be mentioned in the title or summary of an article; b) the fact that the identity of the SNP associated metabolites was not always known; and c) indirect relationships between some eGene products and metabolites. That there is more literature support for eGene-metabolite relationships than for the eSNP-metabolite or eSNP-eGene associations highlights the utility of our approach in bridging the existing knowledge gap.

Our agnostic systems-based approach is helpful to unravel the wholistic genetic architecture that underlies phenotypes including metabolism. Homeostatic metabolite levels in the blood are the sum of the rates of importation from the environment, production and degradation in all of the tissues that contact blood^37–39^. Thus, it is important to include investigations of potential genetic mechanisms in all body tissues. Notably, previous studies have demonstrated the effects of genetic interactions in tissues that seem irrelevant to the phenotypes under investigation^40–43^. We propose that eQTLs affect metabolism by modulating the expression of genes that encode the protein components (*i.e.* enzymes, transcription factors, inhibitors, transporters, receptors *etc*.) of metabolism.

In conclusion, we have shown that blood metabolite-associated SNPs mark expression quantitative trait loci that affect the expression of proximal and distal genes in tissues that are relevant to the associated metabolites. The hundreds of novel eSNP-eGene-metabolite connections identified in this study are a useful resource for further empirical study of the genetic influences of the human metabolome.

## Methods

### Data sources

The 145 metabolite-associated Single Nucleotide Polymorphisms (SNPs) investigated in this study are the genome-wide significant hits from Shin et al^5^. Genomic positions of SNPs were obtained from the human hg19 genome build chromosome bed files downloaded from NCBI (See Data Availability). Gene synonyms and full names were obtained from NCBI’s gene information dataset (ftp.ncbi.nih.gov:gene/DATA/GENE_INFO/Mammalia/Homo_sapiens.gene_info.gz). We used the GENCODE transcript model (See Data Availability) as reference for gene annotations, which is the same reference used in GTEx. All isoforms of a gene were collapsed into a single gene. The human genome used in this study is the hg19 (GRChr37) build of the human genome release 75 (See Data Availability).

### Identification of SNP target genes

The CoDeS3D^24^ pipeline was used to identify target genes of SNPs (Supplementary Figure 1). In summary, restricted fragments that harbor the SNPs were identified in the human genome. Chromatin interaction (Hi-C) libraries from eight cell lines (*i.e.* GM12878, HeLa, HMEC, HUVEC, IMR90, K562, KBM7 and NHEK; Supplementary Table 1)^25^ were then queried to identify gene fragments that are captured as spatially interacting with the SNP-containing fragments. The resulting spatial SNP-gene pairs were used to query GTEx V7 to identify eSNP-eGene pairs in 48 human tissues. eSNP-eGene-tissue associations with adjusted p values < 0.05 (*FDR*, Benjamini-Hochberg correction) were deemed significant.

### Annotation of genes involved in metabolism

The Kyoto Encyclopedia of Genes and Genomes (KEGG) PATHWAY^44^ (https://www.kegg.jp/kegg/pathway.html, accessed on 01/07/2018) database was queried with a list of the eGenes to identify their associated pathways (Supplementary Figure 1). The retrieved results were analyzed to identify their first level pathway maps (*e.g.* metabolism, genetic information processing) and second level maps (*e.g.* carbohydrate metabolism, transcription). See Code Availability for the Python scripts used for this analysis.

### Drug associations and protein classification

Data from The Human Protein Atlas^45^ (https://www.proteinatlas.org/, downloaded on 25/03/19) version 18.1 was queried to obtain the protein classes of eGenes. The Drug Gene Interaction database^46^ (DGIdb) was interrogated for information on drugs that target the gene products (Supplementary Figure 1). The mechanisms of action of the drugs were also obtained. See Code Availability for the Python scripts used for this analysis.

### Literature support for associations

To find evidence for eSNP-eGene, eSNP-metabolite, or eGene-metabolite associations in literature, we performed a text mining algorithm as follows. We employed the Bio. Entrez python API (see Code Availability) to search the underlying MEDLINE database to retrieve the PubMed IDs for articles that contain, in their titles or abstracts, exact matches of 1) eSNPs (rsIDs) and their linked eGenes (or gene synonyms or full names); 2) eSNPs (rsIDs) and at least one of the metabolites associated to the eSNP by Shin et al^5^; or 3) eGenes (or their synonyms or full names) and at least one of the metabolites associated to the corresponding eSNP. We then identified intersecting articles for corresponding eSNP-eGene, eSNP-metabolite, and eGene-metabolite associations. Articles supporting gene-metabolite associations not reported in Shin et al were manually curated by at least two persons.

### URLS

CoDeS3D pipeline: https://github.com/Genome3d/codes3d-v1 The Drug Gene Interaction database: https://dgidb.org

GTEx portal: https://www.gtexportal.org/home/

The KEGG PATHWAY database: https://www.kegg.jp/kegg/pathway.html The Human Protein Atlas: https://www.proteinatlas.org/

### Data and Code Availability

Supplementary tables are available at https://www.doi.org/10.17608/k6.auckland.8116097

Scripts used for data curation, analysis, and visualisation are available at https://github.com/Genome3d/blood-metabolites-regulation.git

Human genome build hg19 (GRChr37) was downloaded from ftp.ensembl.org/pub/release-75/fasta/homo_sapiens/

SNP annotations (human genome, build hg19) were obtained from ftp.ncbi.nih.gov/snp/organisms/human_9606_b146_GRCh37p13/BED

Gene annotations (Transcript model from GENCODE) were downloaded from http://www.gtexportal.org/static/datasets/gtex_analysis_v6p/reference/gencode.v19.genes.v6p_model.patched_contigs.gtf.gz

## Acknowledgements

This work was funded by High Value Nutrition National Science (MBIE/HVN grant #3710040) to JOS & TF. JOS and WS are funded by a Royal Society of New Zealand Marsden Fund [Grant 16-UOO-072].

## Author Contributions

TF ran analyses, wrote scripts, interpreted data and wrote the manuscript. WS contributed to data interpretation and commented on the manuscript. NK and JL contributed to the literature text mining. JI contributed to data interpretation and commented on the manuscript. JOS directed the study, contributed to data interpretation, and commented on the manuscript. JOS is guarantor for this article.

## Supplementary Information

### Supplementary Figures

**Supplementary Figure 1.**
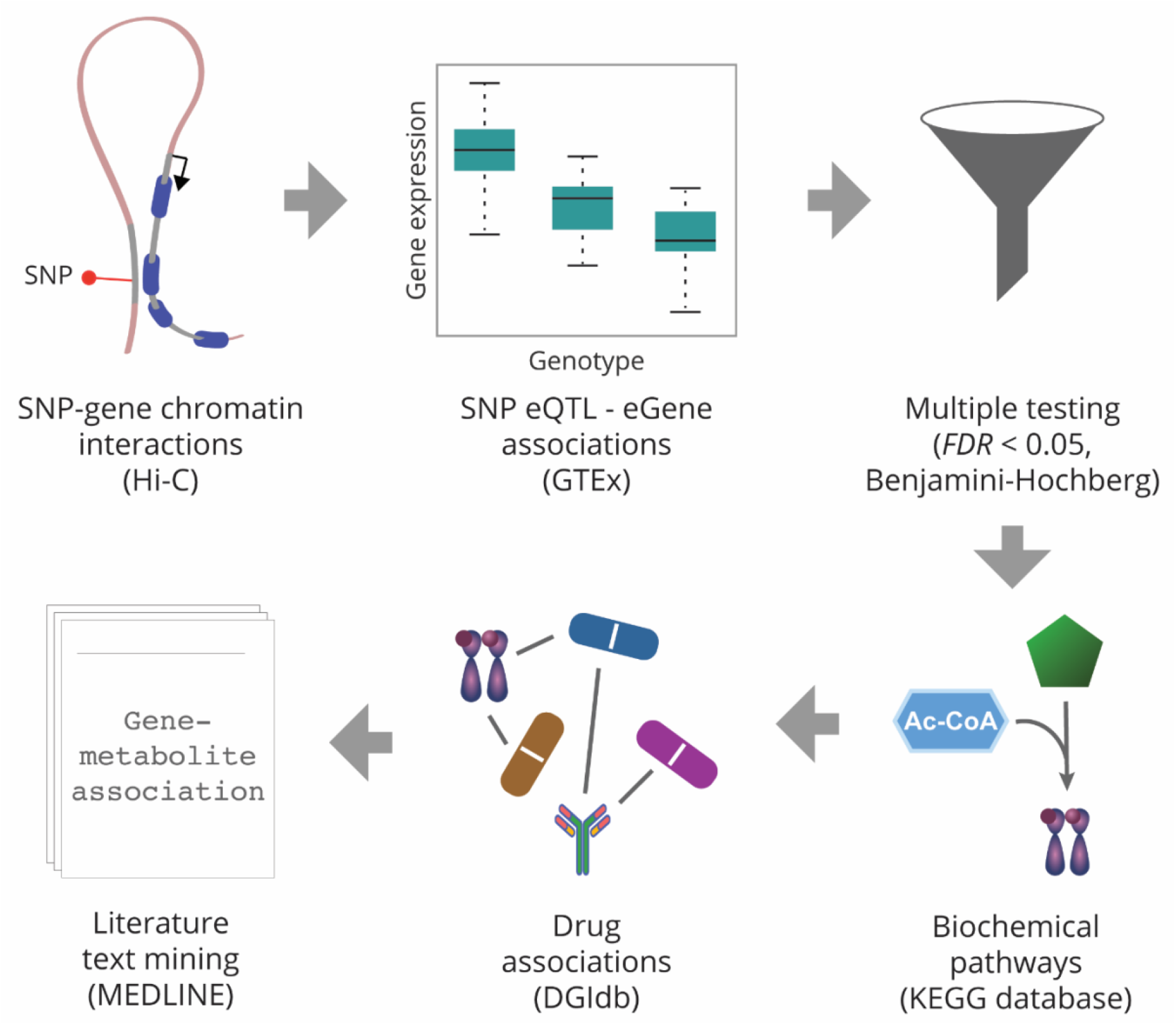
Methods workflow. Genes in restriction fragments that are spatially interacting with fragments containing metabolite-associated SNPs were identified using Hi-C libraries. The resulting spatial SNP-gene pairs were used to query GTEx for tissue eQTL interactions. Only significant (FDR < 0.05) tissue eSNP-eGene associations were further analyzed for ontology, drug associations and literature text mining.

